# The membrane-associated ubiquitin ligase MARCHF8 stabilizes the human papillomavirus oncoprotein E7 by degrading CUL1 and UBE2L3 in head and neck cancer

**DOI:** 10.1101/2023.11.03.565564

**Authors:** Mohamed I. Khalil, Canchai Yang, Lexi Vu, Smriti Chadha, Harrison Nabors, Claire D. James, Iain M. Morgan, Dohun Pyeon

**Author notes:** Address correspondence to Dohun Pyeon, and Mohamed Khalil.

## Abstract

The human papillomavirus (HPV) oncoprotein E7 is a relatively short-lived protein required for HPV-driven cancer development and maintenance. E7 is degraded through ubiquitination mediated by cullin 1 (CUL1) and the ubiquitin-conjugating enzyme E2 L3 (UBE2L3). However, E7 proteins are maintained at high levels in most HPV-positive cancer cells. A previous proteomics study has shown that UBE2L3 and CUL1 protein levels are increased by the knockdown of the E3 ubiquitin ligase membrane-associated ring-CH-type finger 8 (MARCHF8). We have recently demonstrated that HPV upregulates MARCHF8 expression in HPV-positive keratinocytes and head and neck cancer (HPV+ HNC) cells. Here, we report that MARCHF8 stabilizes the E7 protein by degrading the components of the SKP1-CUL1-F-box (SCF) ubiquitin ligase complex in HPV+ HNC cells. We found that *MARCHF8* knockdown in HPV+ HNC cells drastically decreases the E7 protein level while increasing the CUL1 and UBE2L3 protein levels. We further revealed that the MARCHF8 protein binds to and ubiquitinates CUL1 and UBE2L3 proteins and that MARCHF8 knockdown enhances the ubiquitination of the E7 protein. Conversely, the overexpression of CUL1 and UBE2L3 in HPV+ HNC cells decreases E7 protein levels and suppresses tumor growth in vivo. Our findings suggest that HPV-induced MARCHF8 prevents the degradation of the E7 protein in HPV+ HNC cells by ubiquitinating and degrading CUL1 and UBE2L3 proteins.

**IMPORTANCE:** Since HPV oncoprotein E7 is essential for virus replication, HPV has to maintain high levels of E7 expression in HPV-infected cells. However, HPV E7 can be efficiently ubiquitinated by a ubiquitin ligase and degraded by proteasomes in the host cell. Mechanistically, the components of the E3 ubiquitin ligase complex CUL1 and UBE2L3 play an essential role in E7 ubiquitination and degradation. Here, we show that the membrane ubiquitin ligase MARCHF8 induced by HPV E6 stabilizes the E7 protein by degrading CUL1 and UBE2L3 and blocking E7 degradation through proteasomes. MARCHF8 knockout restores CUL1 and UBE2L3 expression, decreasing E7 protein levels and inhibiting the proliferation of HPV-positive cancer cells. Additionally, overexpression of CUL1 or UBE2L3 decreases E7 protein levels and suppresses in vivo tumor growth. Our results suggest that HPV maintains high E7 protein levels in the host cell by inducing MARCHF8, which may be critical for cell proliferation and tumorigenesis.

## INTRODUCTION

Human papillomaviruses (HPVs) are non-enveloped DNA viruses that infect skin keratinocytes (1). The high-risk HPV genotypes such as HPV16 and HPV18 are causally associated with over 95% of cervical cancers, 25% of head and neck cancers (HNC), and most anogenital cancers (2, 3). High-risk HPV E6 and E7 interact with various cellular proteins to modulate almost all oncogenic mechanisms defined as cancer hallmarks (reviewed in 4). Thus, HPV-driven cancers are addicted to E6 and E7 expression, and their survival requires sustaining high E7 expression (5–7).

HPV16 E6 and E7 facilitate the ubiquitination and degradation of various host proteins by interacting with E2 ubiquitin-conjugating enzymes and E3 ubiquitin ligases (8–12). While E6 degrades p53 through E6AP, E7 degrades host proteins such as pocket proteins (pRb, p107, and p130) and the tyrosine-protein phosphatase non-receptor type 14 (PTPN14) through the cullin-RING E3 ubiquitin ligase complex (13–16). On the other hand, the HPV16 E7 protein is also degraded through ubiquitination by the S-phase kinase-associated protein (SKP)-Cullin-F box (SCF) ubiquitin ligase complex containing ubiquitin-conjugating enzyme E2 L3 (UBE2L3, also called UBCH7), cullin 1 (CUL1), and S-phase kinase-associated protein 2 (SKP2) (17). The SCF ubiquitin ligase complex ubiquitinates many cell-cycle-regulatory proteins, such as E2F1, p27, origin recognition complex subunit 1 (ORC1), and cyclin D1 (18–21). UBE2L3 interacts with various E3 ligases containing HECT (homologous of E6AP C-terminus) or RING (Really Interesting New Gene) finger domains (22, 23). Following SCF-mediated ubiquitination, E7 is degraded via the 26S proteasome (17, 24). Despite the efficient degradation mechanism in host cells, HPV-positive (HPV+) cancer cells maintain high levels of E7 proteins. Previous studies have shown that the E7 protein is stabilized by various mechanisms, such as deubiquitination (25) and phosphorylation (26). However, little is known about how the E7 protein overcomes its degradation by the SCF ubiquitin ligase complex mediated ubiquitination in HPV+ cancer cells.

A recent study showed that MARCHF8 knockdown in esophageal squamous cell carcinoma cells increases the levels of CUL1 and UBE2L3 proteins (27). MARCHF8 is a member of the MARCHF family of E3 ubiquitin ligases, which ubiquitinates diverse immune receptors, such as major histocompatibility complexes (MHC-I and MHC-II) (28, 29) and IL-1 receptor accessory protein (IL1RAP) (30). We have recently revealed that MARCHF8 is significantly upregulated in HPV+ HNC cells and inhibits apoptosis by ubiquitinating and degrading the death receptors from the tumor necrosis factor (TNF) receptor superfamily, FAS, and TNF-related apoptosis-inducing ligand receptors 1 and 2 (TRAIL-R1 and TRAIL-R2) (31).

Here, we show that MARCHF8 binds to and ubiquitinates CUL1 and UBE2L3 proteins in HPV+ HNC cells. Furthermore, knockdown or knockout of *MARCHF8* in HPV+ HNC cells significantly increased CUL1 and UBE2L3 protein levels, decreasing the levels of HPV E7 protein. Conversely, overexpression of CUL1 and UBE2L3 in HPV+ HNC cells decreases E7 protein levels and suppresses tumor growth in vivo. These findings suggest that HPV-induced MARCHF8 stabilizes E7 proteins to maintain the high levels of E7 in HPV+ HNC cells.

## RESULTS

### CUL1 and UBE2L3 overexpression increases the ubiquitination and decreases the E7 protein levels in HPV+ HNC cells

HPV E7 was previously shown to be ubiquitinated and degraded by CUL1 and UBE2L3 in cervical cancer cells (17) (**Fig. 1A**). Hence, to examine if CUL1 and UBE2L3 enhance ubiquitination and degradation of HPV16 E7 protein in the HPV+ HNC cells, we generated SCC152 and SCC2 cells overexpressing CUL1 or UBE2L3 using lentiviral transduction and blasticidin selection. Western blotting showed that *CUL1* or *UBE2L3* overexpression significantly decreased E7 protein levels in both SCC152 (**Fig. 1B and 1C**) and SCC2 (**Fig. 1D and 1E**) cells. These results confirm that CUL1 and UBE2L3 reduce E7 protein levels. Interestingly, CUL1 overexpression increased UBE2L3 protein levels in SCC152 and SCC2, and UBE2L3 overexpression increased CUL1 protein levels in the two cell lines. Next, to determine if CUL1 and UBE2L3 overexpression enhances the ubiquitination of E7 proteins, we pulled down ubiquitinated proteins in whole cell lysates from the proteasome inhibitor MG132-treated SCC152 cells overexpressing CUL1 or UBE2L3 using anti-ubiquitin antibody-conjugated magnetic beads. The results showed that ubiquitination levels of E7 (**Fig. 1F**) were increased by CUL1 and UBE2L3 overexpression in SCC152 cells despite the significantly lower levels of input E7 proteins compared to SCC152 cells with vector. These results suggest that CUL1 and UBE2L3 enhance E7 ubiquitination and degradation.

**Fig. 1.**
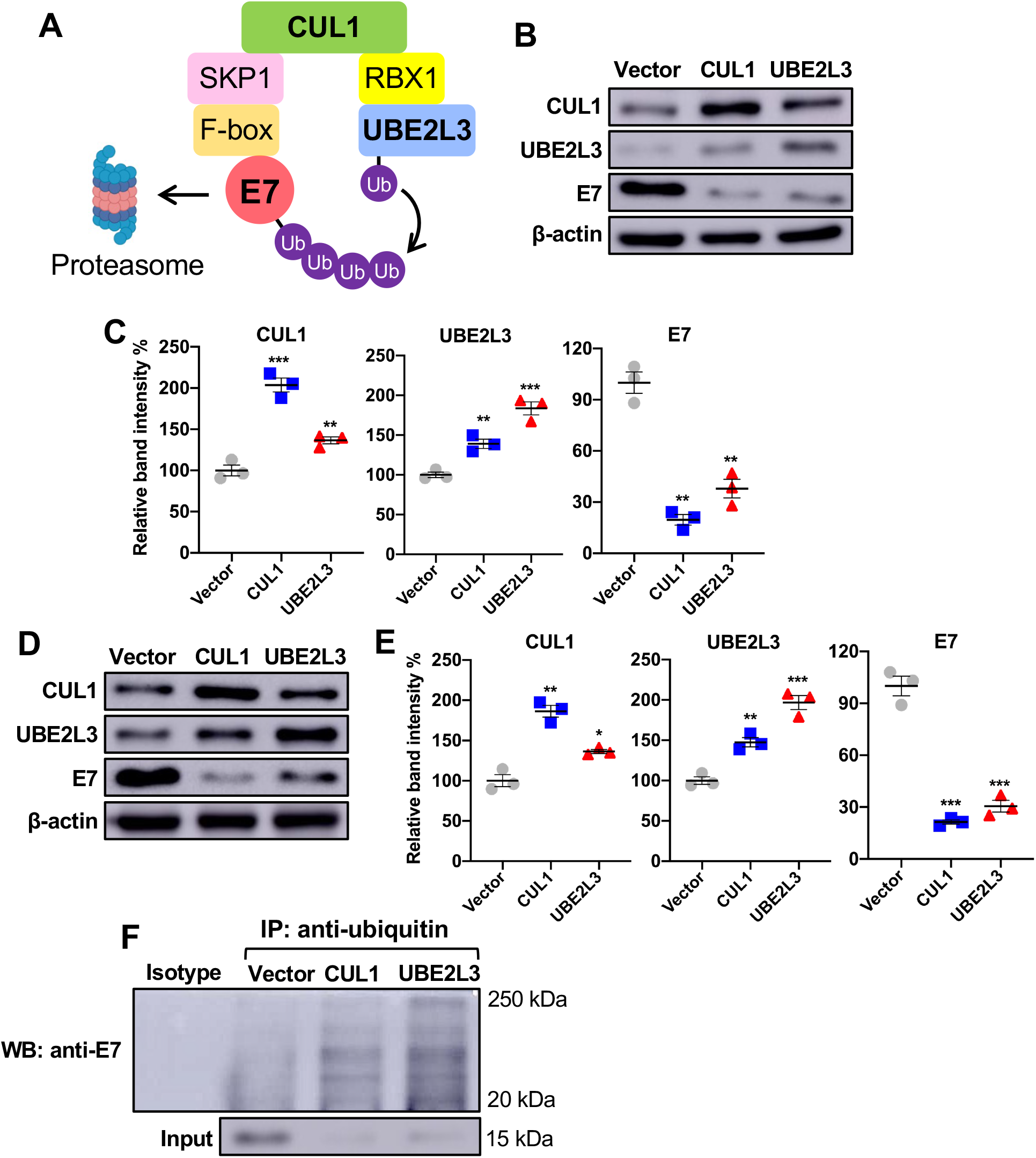
CUL1 and UBE2L3 overexpression increases the ubiquitination and decreases HPV16 E7 protein level in HPV+ HNC cells. (**A**) The schematic model of E7 degradation through CUL1 and UBE2L3-mediated ubiquitination. CUL1 or UBE2L3 is overexpressed in HPV+ HNC (SCC152) (**B-C**) and SCC2 (**D-E**) cells using lentiviral transduction. CUL1, UBE2L3, HPV16 E7, and MARCHF8 proteins were detected by western blotting. The relative band intensities were quantified using NIH ImageJ in SCC152 (**C**) and SCC2 (**E**). β-actin was used as an internal control. (**F**) SCC152 cells with CUL1 or UBE2L3 overexpression were treated with MG132 (10 μM). Ubiquitinated proteins were pulled down from the cell lysate using anti-ubiquitin antibody-conjugated magnetic beads, and E7 protein was detected by western blotting. The data shown are means ± SD of three independent experiments. Student’s t-test was used to determine *p* values. \**p* < 0.05, \*\**p* < 0.01, \*\*\**p* < 0.001.

### CUL1 and UBE2L3 expression is downregulated in HPV+ HNC cells

To determine if HPV alters the endogenous CUL1 and UBE2L3 expression, we measured CUL1 and UBE2L3 protein levels in HPV+ HNC cells (SCC2, SCC90, and SCC152) compared to HPV-HNC cells (SCC1, SCC9, and SCC19) and normal keratinocytes (N/Tert-1) by western blotting. The results showed that CUL1 and UBE2L3 protein levels are significantly lower in HPV+ HNC cells compared to HPV-HNC cells and normal keratinocytes (**Fig. 2A and 2B**). To examine if CUL1 and UBE2L3 expression is transcriptionally regulated in HPV+ HNC cells, we measured CUL1 and UBE2L3 mRNA levels in HPV+ and HPV-HNC cells along with N/Tert-1 cells by RT-qPCR. Our results showed that CUL1 mRNA expression was not significantly changed in SCC90 and SCC152 cells but upregulated in SCC2 cells compared to N/Tert-1 cells (**Fig. 2C**). On the other hand, UBE2L3 mRNA expression was significantly upregulated in all HPV+ HNC cells but downregulated in all HPV-HNC cells compared to N/Tert-1 cells (**Fig. 2C**). These results suggest that the low CUL1 and UBE2L3 protein levels in HPV+ HNC cells are not caused by transcriptional regulation.

**Fig. 2.**
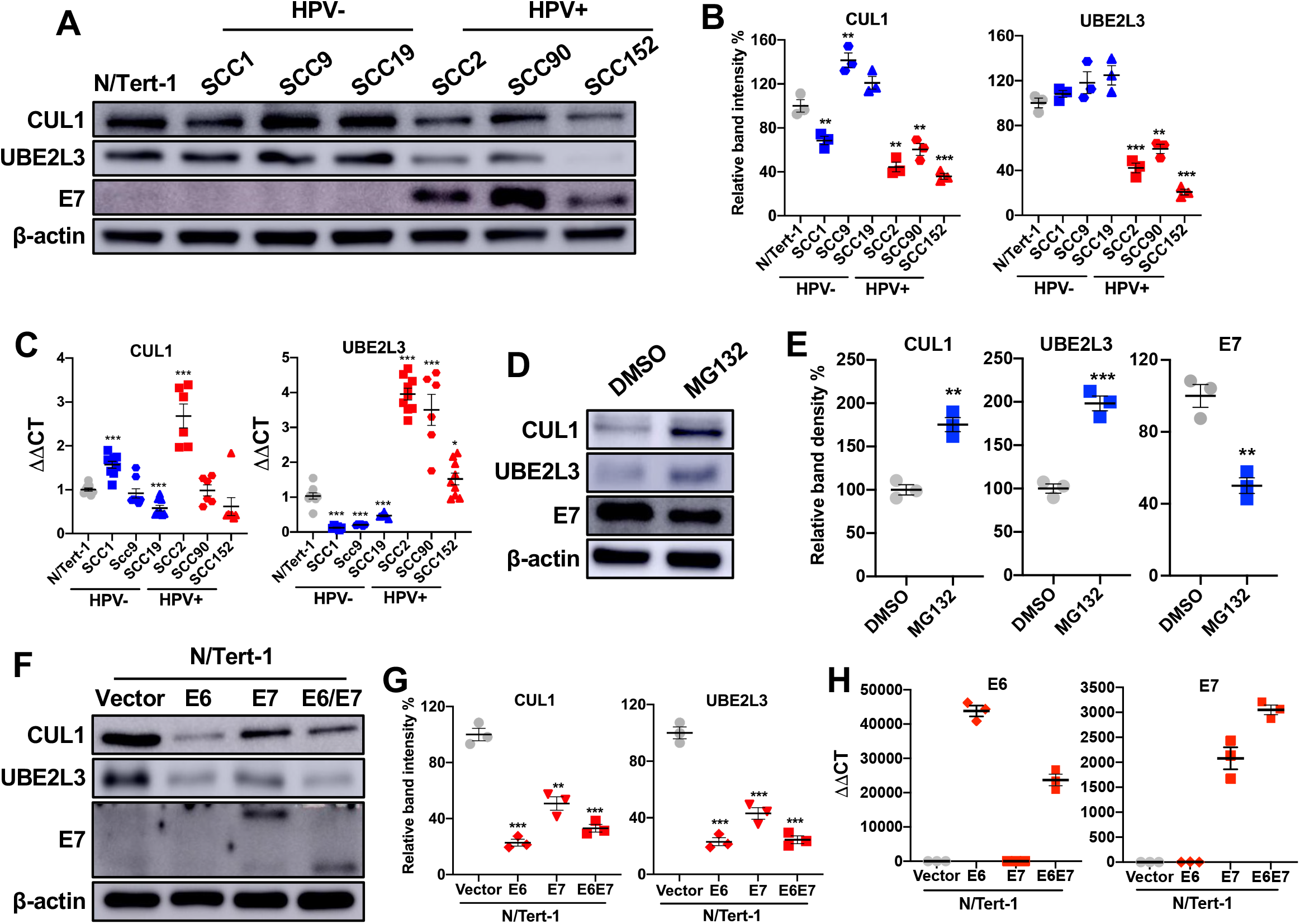
CUL1 and UBE2L3 protein levels are low in HPV+ HNC cells. CUL1 and UBE2L3 protein levels in normal (N/Tert-1), HPV-HNC (SCC1, SCC9, and SCC19), and HPV+ HNC (SCC2, SCC90, and SCC152) cells were determined by western blotting (**A**). Relative band intensities were quantified using NIH ImageJ (**B**). HPV16 E7 and β-actin were used as viral and internal controls, respectively. CUL1 and UBE2L3 mRNA expression levels in normal (N/Tert-1), HPV+ HNC (SCC-2, SCC-90, and SCC-152), and HPV-HNC (SCC-1, SCC-9, and SCC-19) cells were quantified by RT-qPCR (**C**). The data shown are normalized by the GAPDH mRNA level as an internal control. CUL1, UBE2L3, and E7 protein levels in SCC152 cells treated with MG132 (10 μM) or DMSO (**D**). Relative band intensities were quantified using NIH ImageJ (**E**). CUL1 and UBE2L3 proteins were detected in N/Tert-1 cells expressing HPV16 E6, E7, or E6 and E7 by western blotting (**F**). An HA tag increases the size of HPV16 E7 in N/Tert-1 E7 cells to ∼22 kDa from ∼17 kDa. Relative band intensities were quantified using NIH ImageJ (**G**). Total RNA was extracted from N/Tert-1 containing an empty vector and N/Tert-1 cells expressing HPV16 E6, E7, or E6 and E7 (E6E7). The HPV16 E6 and E7 mRNA expression levels were quantified by RT-qPCR (**H**). The data shown are normalized by the GAPDH mRNA level as an internal control. All experiments were repeated at least three times, and the data shown are means ± SD of three independent experiments. Student’s t-test determined *p* values. \**p* < 0.05, \*\**p* < 0.01, \*\*\**p* < 0.001.

To determine if proteasome-dependent protein degradation causes the low CUL1 and UBE2L3 protein levels in HPV+ HNC cells, CUL1 and UBE2L3 protein levels were measured in MG132-treated SCC152 cells. The results showed that MG132 treatment increased the CUL1 and UBE2L3 protein levels in SCC152 cells (**Fig. 2D and 2E**). These results suggest that the proteasome-dependent degradation of CUL1 and UBE2L3 proteins is enhanced in HPV+ HNC cells. Surprisingly, the E7 protein level was not increased by MG132 treatment in the SCC152 cells (**Fig. 2D and 2E**). This may be due to the significantly increased CUL1 and UBE2L3 proteins by MG132, which enhance E7 ubiquitination, compensating for proteasome inhibition.

Next, to assess if the HPV oncoproteins E6 and/or E7 are responsible for the low CUL1 and UBE2L3 protein levels, we measured CUL1 and UBE2L3 protein levels in N/Tert-1 cells expressing HPV16 E6 (N/Tert-1 E6), E7 (N/Tert-1 E7), and E6E7 (N/Tert-1 E6E7) compared to control N/Tert-1 cells containing an empty vector (N/Tert-1 vector) by western blotting. The HPV16 E6 and E7 mRNA (**Fig. 2H**) and E7 protein levels (**Fig. 2F**) in these N/Tert-1 cells were verified by RT-qPCR and western blotting, respectively. The results showed that the protein levels of both CUL1 and UBE2L3 were significantly decreased in HPV16 E6, HPV16 E7, and HPV16 E6E7 cells compared to N/Tert-1 vector cells (**Fig. 2F and 2G**). These results suggest that either E6 or E7 expression is sufficient for decreasing CUL1 and UBE2L3 protein levels in normal keratinocytes.

### MARCHF8 knockdown increases the CUL1 and UBE2L3 protein levels and decreases the E7 protein level in HPV+ HNC cells

**The** E3 ubiquitin ligase MARCHF8 ubiquitinates several cellular proteins for degradation. By analyzing proteomics data from a previous study in esophageal squamous cell carcinoma (27), we found that the CUL1 and UBE2L3 proteins are among the cellular proteins increased by MARCHF8 knockdown. Thus, we hypothesized that MARCHF8 plays an essential role in maintaining high E7 protein levels in HPV+ HNC cells by degrading CUL1 and UBE2L3 proteins. To test the hypothesis, we knocked down MARCHF8 expression in two HPV+ HNC cells, SCC152 and SCC2, using multiple shRNAs against MARCHF8 (shR-MARCHF8) delivered by lentiviral transduction. All SCC152 (**Fig. 3A and 3C**) and SCC2 (**Fig. 3B and 3D**) cell lines with shR-MARCHF8 showed at least a 50% decrease in MARCHF8 protein levels compared to the cells with scrambled shRNA (shR-scr). As hypothesized, western blotting showed that CUL1 and UBE2L3 protein levels are significantly increased by MARCHF8 knockdown (**Fig. 3A - 3D**). Interestingly, HPV16 E7 protein levels are dramatically decreased by MARCHF8 knockdown, showing an inverse correlation with the CUL1 and UBE2L3 protein levels (**Fig. 3A - 3D**). The decreased E7 protein levels are also validated by the significant increase in pRb protein levels by MARCHF8 knockdown (**Fig. 3A - 3D**). To test if MARCHF8 knockdown affects mRNA expression of CUL1 and UBE2L3, we measured mRNA levels of CUL1 and UBE2L3 in SCC152 and SCC2 cells with MARCHF8 knockdown by RT-qPCR. The results showed a significant decrease in CUL1 and UBE2L3 mRNA levels by MARCHF8 knockdown in SCC152 cells (**Fig. 3E**) but not in SCC2 cells (**Fig. 3F**). These results indicate that the increase of the CUL1 and UBE2L3 protein levels by MARCHF8 knockdown in HPV+ HNC cells is independent of CUL1 and UBE2L3 mRNA expression. We also observed a slight decrease in HPV16 E7 mRNA levels by MARCHF8 knockdown (**Fig. 3E and 3F**). However, as the E7 mRNA levels are still high despite the MARCHF8 knockdown, it is unlikely that the slight decrease of E7 mRNA levels entirely causes the considerable reduction of the E7 protein levels by the MARCHF8 knockdown. Thus, our findings suggest that HPV-induced MARCHF8 is responsible for the low CUL1 and UBE2L3 and high E7 protein levels in HPV+ HNC cells.

**Fig. 3.**
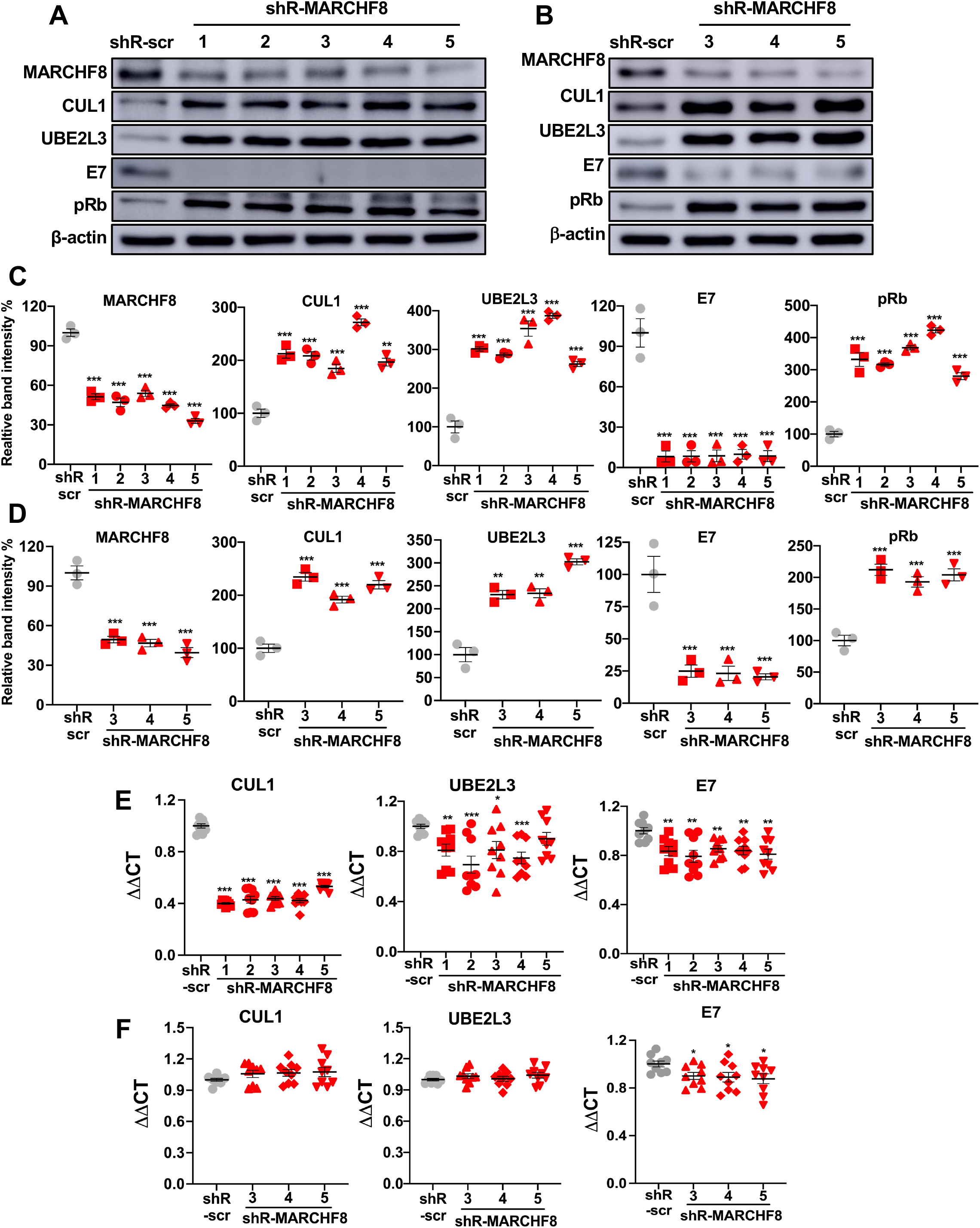
Knockdown of MARCHF8 expression increases CUL1 and UBE2L3 protein levels and decreases HPV16 E7 protein levels in HPV+ HNC cells. SCC152 (**A & C**) and SCC2 (**B & D**) cells were transduced with five and three lentiviral shRNAs against MARCHF8 (shR-MARCHF8), respectively, or scrambled shRNA (shR-scr) as a control. MARCHF8, CUL1, UBE2L3, HPV16 E7, and pRb proteins in SCC152 (**A**) and SCC2 (**B**) were detected by western blotting. The relative band intensities were quantified using NIH ImageJ (**C** and **D**). β-actin was used as an internal control. The mRNA levels of CUL1, UBE2L3, and E7 were quantified by RT-qPCR in SCC152 (**E**) and SCC2 (**F**) cells transduced with five and three lentiviral shRNAs against MARCHF8 (shR-MARCHF8), respectively, or scrambled shRNA (shR-scr) as a control. The data shown are normalized by the GAPDH mRNA level as an internal control. The data shown are means ± SD of three independent experiments. Student’s t-test determined *p* values. \**p* < 0.05, \*\**p* < 0.01, \*\*\**p* < 0.001.

### MARCHF8 interacts with and ubiquitinates CUL1 and UBE2L3 proteins in HPV+ HNC cells

As MARCHF8 knockdown increased CUL1 and UBE2L3 protein levels (**Fig. 3A and 3B**), we hypothesized that HPV-induced MARCHF8 ubiquitinates CUL1 and UBE2L3 proteins for proteasomal degradation. First, to determine if MARCHF8 binds to CUL1 and UBE2L3, we pulled down MARCHF8 protein in whole cell lysates from SCC152 cells treated with MG132 using anti-MARCHF8 antibody-conjugated magnetic beads. We detected CUL1 and UBE2L3 proteins by western blotting. The results showed that CUL1 and UBE2L3 proteins were co-immunoprecipitated with MARCHF8 protein (**Fig. 4A**). In contrast, E7 protein was not co-immunoprecipitated with MARCHF8-protein (**Fig. 4A**), indicating that E7 may not be a direct target of MARCHF8. Reciprocally, we pulled down CUL1 or UBE2L3 proteins in the same whole cell lysates from MG132-treated SCC152 cells using anti-CUL1 or anti-UBE2L3 antibodies, respectively and found that MARCHF8 protein is co-immunoprecipitated with CUL1 and UBE2L3 proteins (**Fig. 4B and 4C**). These results suggest that the MARCHF8 protein interacts with CUL1 and UBE2L3 but not the E7 protein.

**Fig. 4.**
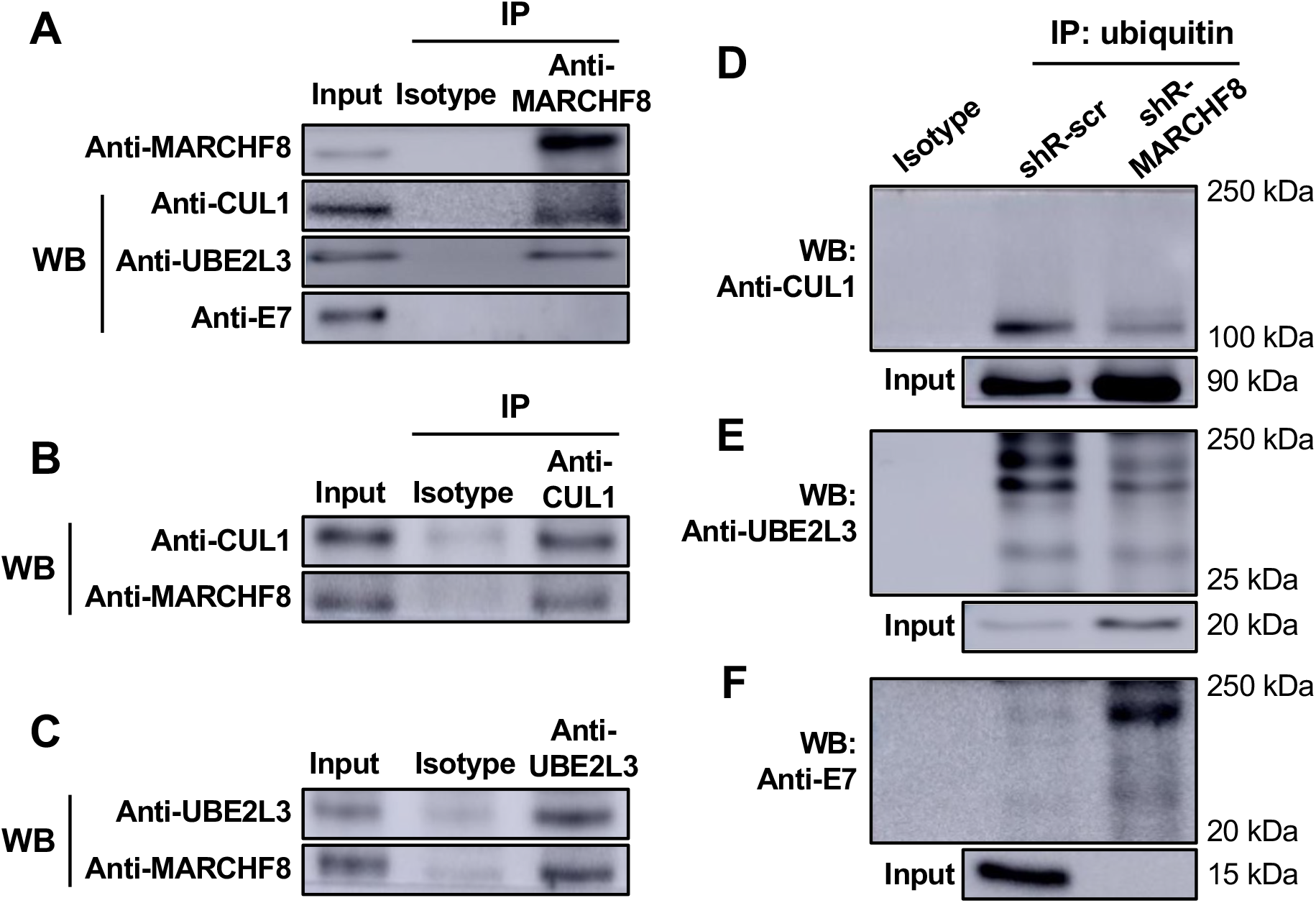
MARCHF8 protein interacts with and ubiquitinates CUL1 and UBE2L3 proteins. MARCHF8 (**A**), CUL1 (**B**), and UBE2L3 (**C**) were pulled down from the lysate of SCC-152 cells treated with MG132 (10 μM) using anti-MARCHF8 (**A**), anti-CUL1 (**B**), and anti-UBE2L3 (**C**) antibodies, respectively. CUL1, UBE2L3, E7, and MARCHF8 were detected from the immunoprecipitated proteins by western blotting. (**D-F**) Ubiquitinated proteins were pulled down from the lysate of SCC152 cells with scrambled shRNA (shR-scr) or shRNA against MARCHF8 (shR-MARCHF8 clone 3) treated with MG132 (10 μM) using anti-ubiquitin antibody-conjugated magnetic beads. CUL1 (**D**), UBE2L3 (**E**), and HPV16 E7 (**F**) proteins were detected in the immunoprecipitated proteins by western blotting. All experiments were repeated at least three times. The data shown are means ± SD of three independent experiments. Student’s t-test determined *p* values. \**p* < 0.05, \*\**p* < 0.01, \*\*\**p* < 0.001.

Next, to determine if MARCHF8 ubiquitinates CUL1 and UBE2L3 proteins, we pulled down ubiquitinated proteins in whole cell lysates from MG132-treated SCC152 cells using anti-ubiquitin antibody-conjugated magnetic beads. The results showed that ubiquitinated forms of CUL1 (**Fig. 4D**) and UBE2L3 (**Fig. 4E**) proteins were decreased in SCC152 cells with MARCHF8 knockdown, despite the significantly higher levels of total input CUL1 and UBE2L3 proteins compared to SCC152 cells with shR-scr (**Fig. 4D and 4E**). These results suggest that the decrease of CUL1 and UBE2L3 protein levels in HPV+ HNC cells is likely mediated by MARCHF8-driven ubiquitination and protein degradation. To determine if the ubiquitination of E7 protein is increased by MARCHF8 knockdown, we detected HPV16 E7 protein from the same pulldowns of ubiquitinated proteins. We found that the levels of ubiquitinated E7 protein were dramatically increased by MARCHF8 knockdown, showing an inverse correlation with the ubiquitination levels of CUL1 and UBE2L3 proteins (**Fig. 4F**). Together, these results suggest that MARCHF8 binds to and ubiquitinates CUL1 and UBE2L3 to prevent ubiquitination of E7 protein.

### MARCHF8 knockdown suppresses HPV+ HNC cell proliferation

It is well known that the oncoprotein E7 promotes cell cycle progression and proliferation of HPV-infected cells (32, 33). Thus, we tested if the decrease of E7 protein levels by MARCHF8 knockdown inhibits the proliferation of SCC152 and SCC2 cells by counting cell numbers over time. The results showed that MARCHF8 knockdown significantly decreases the cell numbers of SCC152 (**Fig. 5A**) and SCC2 (**Fig. 5B**) cells compared to the corresponding control cells with shR-scr. In addition, to determine if MARCHF8 knockdown inhibits the cell cycle progression of HPV+ HNC cells, we stained SCC152 and SCC2 cells with bromodeoxyuridine (BrdU) and the Zombie NIR for measuring de novo DNA synthesis and cell viability, respectively, and analyzed labeled cells by flow cytometry. The results showed that BrdU incorporation in SCC152 and SCC2 cells is significantly decreased by MARCHF8 knockdown (**Fig. 5C and 5D**). These results suggest that MARCHF8 knockdown decreases the proliferation of HPV+ HNC cells, likely due to the reduction of E7 protein levels.

**Fig. 5.**
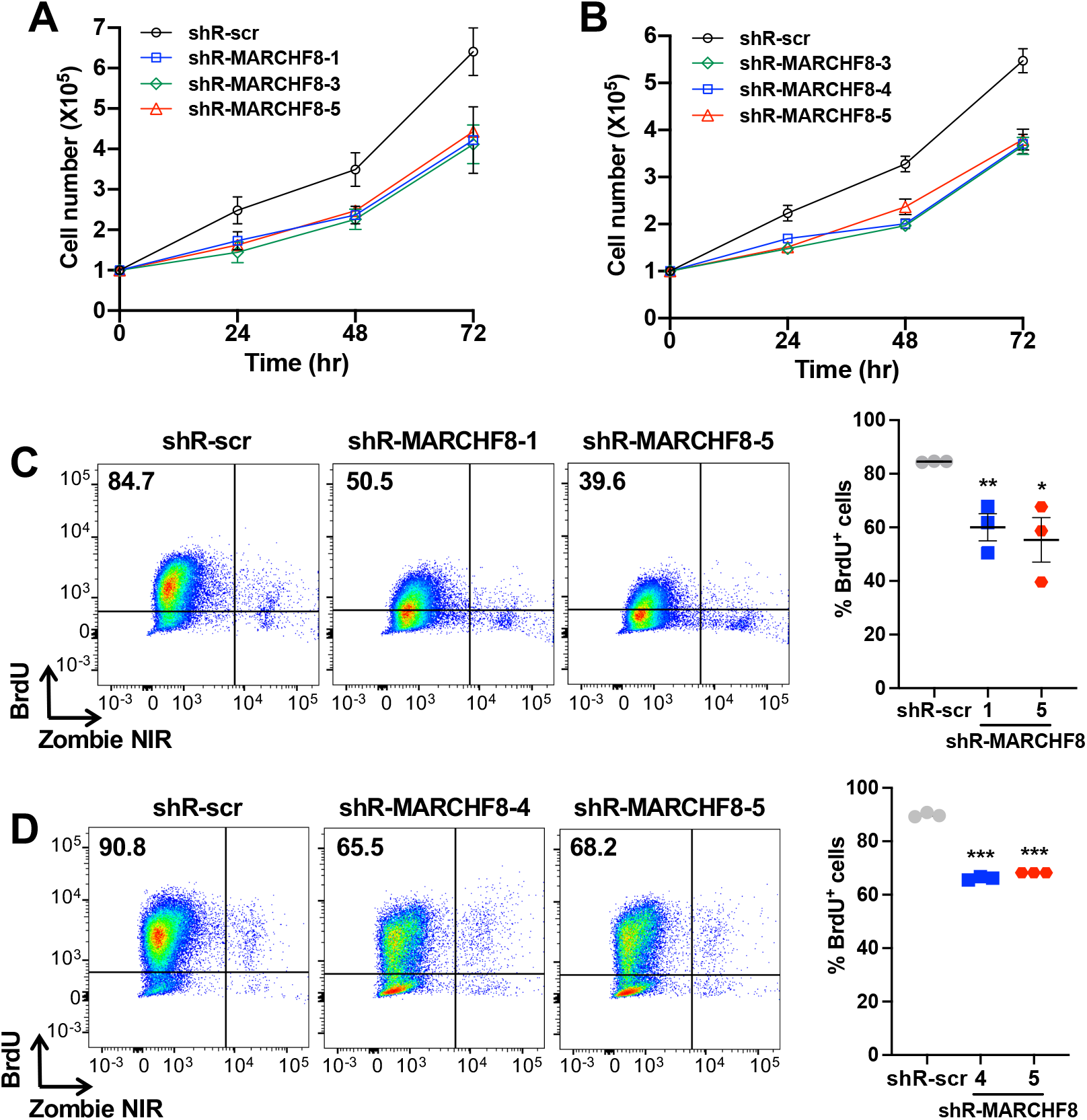
Knockdown of MARCHF8 expression suppresses proliferation of HPV+ HNC cells. Cell proliferation rate was determined by cell counting using two HPV+ HNC cell lines, SCC152 (**A**) and SCC2 (**B**), with scrambled shRNA (shR-scr) or three shRNAs against MARCHF8 (shR-MARCHF8). 1 × 10^5^ cells per well were seeded in a 6-well plate. The cells were detached and counted at 24, 48, and 72 hours. BrdU incorporation assays were performed with SCC152 (shR-MARCHF8 clones 1 and 5) (**C**) and SCC2 (shR-MARCHF8 clones 4 and 5) (**D**) cells with MARCHF8 knockdown. 1 × 10^6^ cells were seeded in 10 cm Petri dishes and incubated for 16 hours with BrdU (10 mM). BrdU incorporation was analyzed by flow cytometry. Cell viability was determined by Zombie NIR staining as a control. All experiments were repeated at least three times, and the data shown are means ± SD. Student’s t-test determined *p* values. \*\*\**p* < 0.001.

### *Marchf8* knockout in HPV+ mouse oral cancer cells restores CUL1 and UBE2L3 protein levels and decreases HPV16 E7 protein levels

To investigate if the degradation of HPV16 E7 protein by CUL1 and UBE2L3 inhibits tumor growth, we utilized a C57BL/6J (B6) mouse oral epithelial (MOE) cell line expressing HPV16 E6/E7 and *Hras* (mEERL) which forms tumors in immunocompetent syngeneic B6 mice (34). First, the protein levels of MARCHF8, CUL1, and UBE2L3 in mEERL cells were determined by comparing them to normal immortalized MOE cells (NiMOE) and mouse HPV-MOE cells transformed with *Hras* and shR-Ptpn14 (MOE/shPtpn14). Consistent with those from human HNC cells, the results showed significantly higher MARCHF8 and lower CUL1 and UBE2L3 protein levels in mEERL cells compared to NiMOE cells and HPV-MOE cells (**Fig. 6A and 6B**). These results suggest that mEERL cells recapitulate our findings from human HPV+ HNC cells, showing that HPV-induced MARCHF8 degrades the CUL1 and UBE2L3 proteins.

**Fig. 6.**
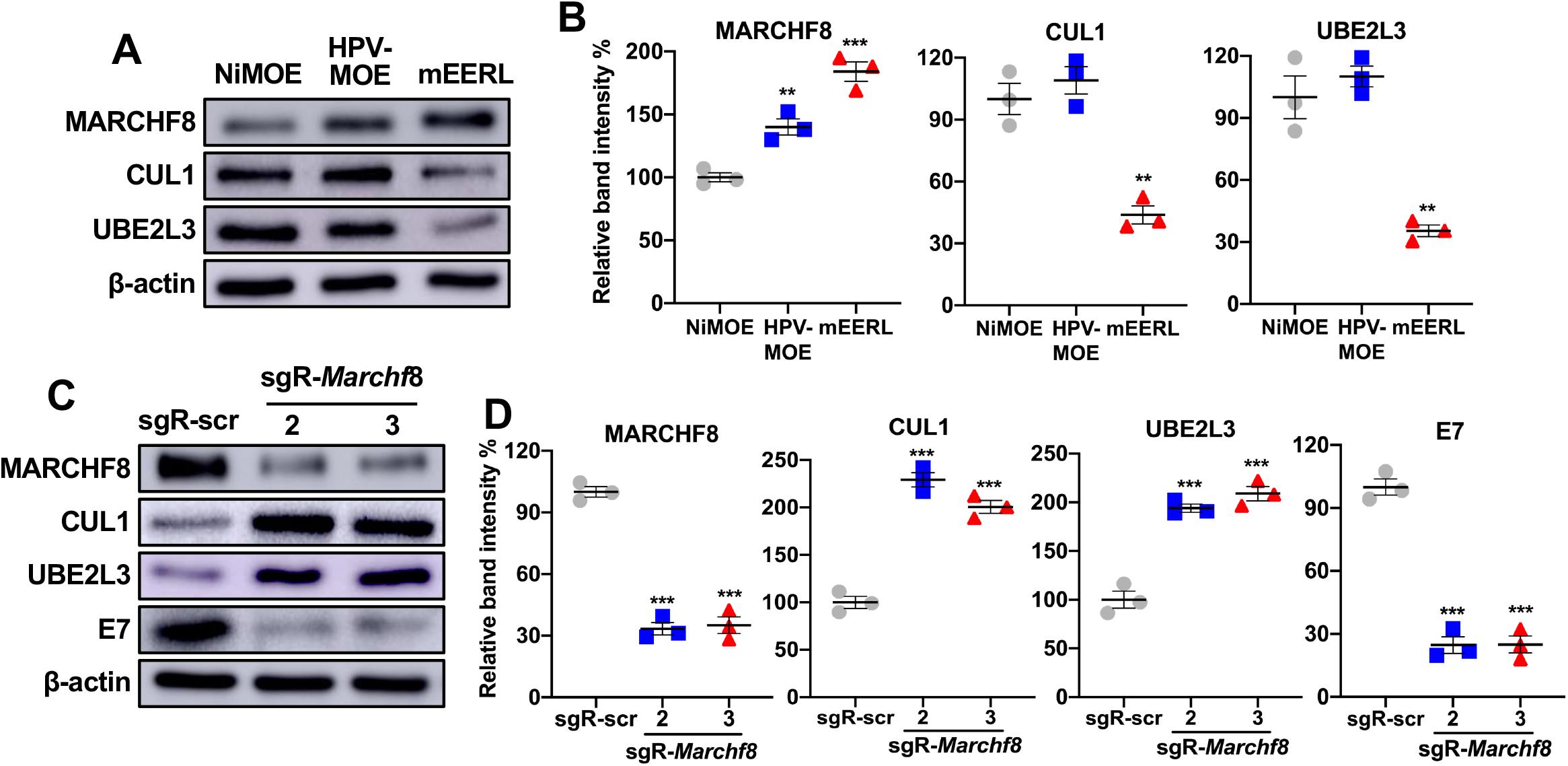
*Marchf8* knockout restores CUL1 and UBE2L3 protein levels in HPV+ mouse oral cancer cells. CUL1 and UBE2L3 protein levels in mouse normal immortalized (NiMOE), HPV-transformed (HPV-MOE), and HPV+ transformed (mEERL) oral epithelial cells were determined by western blotting (**A**). β-actin was used as an internal control. The relative band intensities were quantified using NIH ImageJ (**B**). mEERL cells were transduced with lentiviruses containing Cas9 and one of two sgRNAs against Marchf8 (sgR-*Marchf*8-2 and sgR-*Marchf*8-3) or scrambled sgRNA (sgR-scr). MARCHF8, CUL1, UBE2L3, and HPV16 E7 proteins were detected by western blotting (**C**). Relative band intensities were quantified using NIH ImageJ (**D**). The data shown are means ± SD of three independent experiments. All experiments were repeated at least three times, and the data shown are means ± SD. Student’s t-test determined *p* values. \**p* < 0.05, \*\*\**p* < 0.001.

Next, to determine if high MARCHF8 protein levels in mEERL cells are responsible for the low CUL1 and UBE2L3 protein levels, we measured CUL1 and UBE2L3 protein levels in *Marchf8* knockout mEERL cell lines (mEERL/*Marchf8^-/-^*) previously established using CRISPR/Cas9 (31). Western blotting confirmed that mEERL/*Marchf8*^-/-^ cells have significantly increased CUL1 and UBE2L3 protein levels and decreased HPV16 E7 protein levels compared to mEERL cells with the scrambled control sgRNA (mEERL/scr) (**Fig. 6C and 6D**). These results are consistent with human HPV+ HNC cells presented in **Figs. 3**, suggesting that HPV-induced increase of MARCHF8 proteins in HPV+ HNC cells stabilizes E7 protein by degrading CUL1 and UBE2L3.

### *Cul1* and *Ube2l3* overexpression in HPV+ mouse oral cancer cells delays tumor growth in vivo

To determine whether *Cul1 and Ube2l3* overexpression suppresses HPV+ HNC tumor growth, we generated mEERL cells overexpressing *Cul1* (mEERL/*Cul1*) or *Ube2l3* (mEERL/*Ube2l3*) using lentiviruses and blasticidin selection. Consistent with the results from human HPV+ HNC cells (**Fig. 1B-1E**), *Cul1* or *Ube2l3* overexpression decreased the HPV16 E7 protein level in mEERL cells (**Fig. 7A and 7B**). Next, ten B6 mice were subcutaneously injected with 5 × 10^5^ of either mEERL/*Cul1,* mEERL/*Ube2l3,* or mEERL/scr cells. The results show that all ten mice injected with mEERL/Scr cells vigorously grew tumors (**Fig. 7C and 7F**) and succumbed to tumor burden within ∼7 weeks post-injection (**Fig. 7G**). In contrast, the mice injected with mEERL/*Cul1 or* mEERL/*Ube2l3* cells displayed delayed tumor formation and death by tumor burden (**Fig. 7D and 7G**). Our results suggest that *Cul1* or *Ube2l3* overexpression results in E7 protein degradation and tumor suppression.

**Fig. 7.**
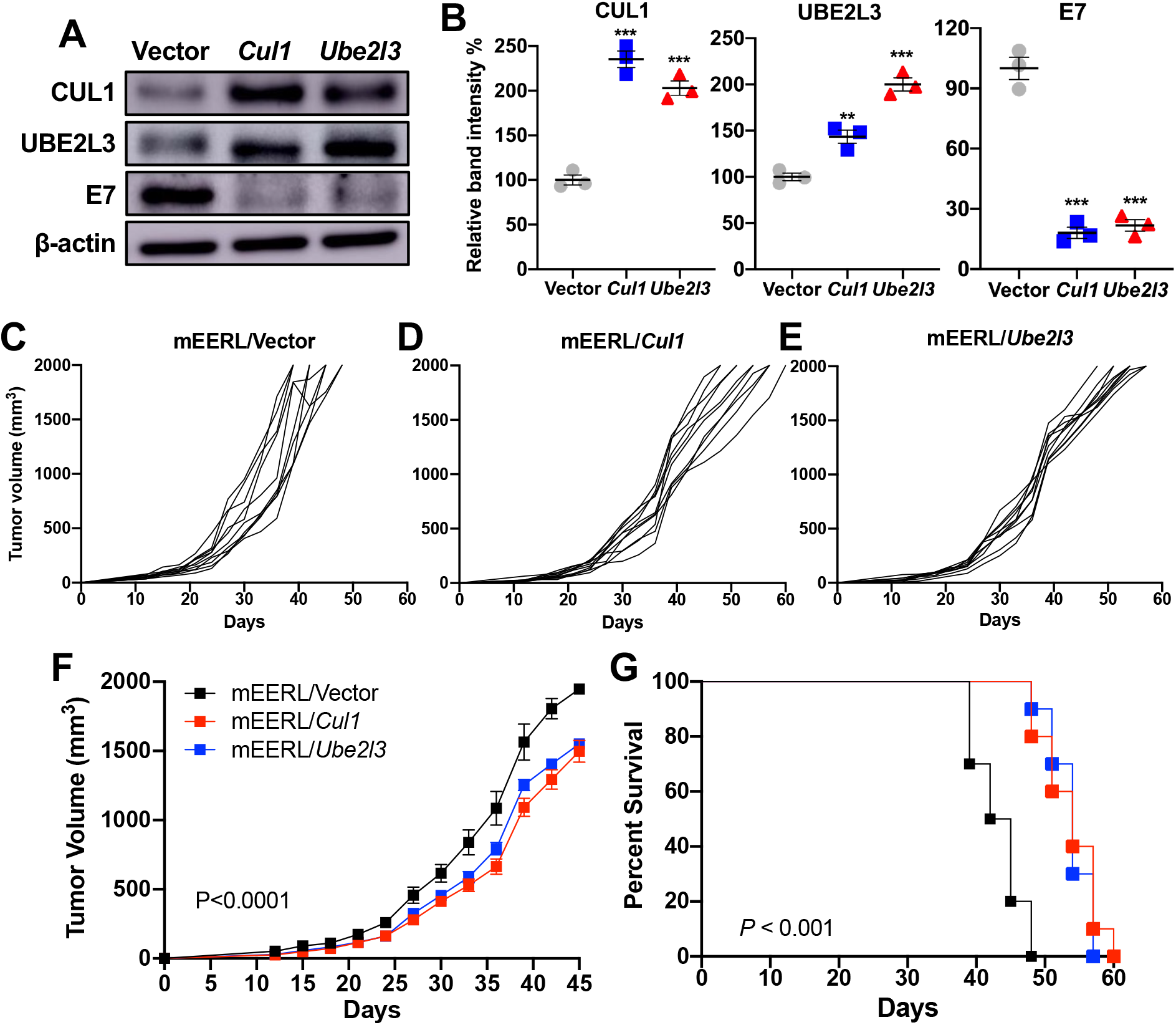
CUL1 and UBE2L3 overexpression suppresses HPV+ HNC tumor growth in vivo. mEERL cells overexpressing *Cul1* or *Ube2l3* were generated by lentiviral transduction of the *Cul1* or *Ube2l3* genes, respectively, and blasticidin selection. CUL1, UBE2L3, and HPV16 E7 proteins were detected by western blotting (**A**). Relative band intensities were quantified using NIH ImageJ (**B**). β-actin was used as an internal control. The data shown are means ± SD of three independent experiments. Student’s t-test determined *p* values. \**p* < 0.05, \*\**p* < 0.01, \*\*\**p* < 0.001. mEERL/vector (**C**), mEERL/*Cul1* (**D**), or mEERL/*Ube2l3* (**E**) cells were injected into the rear right flank of C57BL/6J mice (*n* = 10 per group). Tumor volume was measured twice a week (**C**-**F**). Survival rates of mice were analyzed using a Kaplan-Meier estimator (**G**). The time to event was determined for each group, with the event defined as a tumor size larger than 2000 mm^3^. The data shown are means ± SD. *P* values of mice injected with mEERL/*Cul1 and* mEERL/*Ube2l3* cells compared with mice injected with mEERL/vector cells were determined for tumor growth (**F**) and survival (**G**) by two-way ANOVA analysis. Shown are representative of two independent experiments.

## DISCUSSION

The HPV oncoprotein E7 is essential for virus replication (35), and its continuous expression at high levels is required for HPV-associated cancer progression and maintenance (5, 6, 36, 37). HPV16 E7 interacts with several E3 ubiquitin ligases to facilitate carcinogenesis. It is well understood that E7 induces the proteasomal degradation of various tumor suppressors, such as pRb and mediator of cell motility 1 (MEMO1), by interacting with the CUL2 ubiquitin ligase complex (13, 16, 38, 39). In contrast, E7 also stabilizes host proteins such as the apolipoprotein B mRNA editing enzyme catalytic polypeptide-like 3 (APOBEC3A), a hypoxia-inducible factor (HIF-1a), p53, and p21 (12, 40–42). The dual functions of E7 in host protein degradation suggest intertwined interactions between E7 and various ubiquitin ligases in host cells.

Despite its essential role in cancer progression and maintenance, the E7 protein can be efficiently ubiquitinated and degraded by host factors. E7 protein has a relatively short half-life of less than one hour (43, 44). Mechanistically, the SCF (Skp-Cullin-F box) ubiquitin ligase complex containing CUL1 and UBE2L3 mediates E7 ubiquitination and degradation (17). Although HPV16 E7 contains two lysine residues in positions 60 and 97 (44), the 11 amino acids at the N-terminal residues are essential for ubiquitination and degradation (44). Substituting the two lysines K60 and K97 to arginine in the E7 protein did not abrogate its ubiquitination. In contrast, deleting the N-terminal 11 amino acids prevented E7 ubiquitination and stabilized E7 proteins (44). Mechanistically, the free N-terminal residue of E7 first linearly ubiquitinated, then attaching other ubiquitin groups to the internal lysine of the first ubiquitin moiety at the N-terminal residue (44). The lysine-independent ubiquitination of E7 proteins is similar to the mechanism by which MyoD (45) and Epstein Barr Virus (EBV), Latent Membrane Protein 1 (LMP1) (46) are ubiquitinated (44). To maintain the high E7 levels in HPV+ cancer cells, E7 proteins are stabilized through several mechanisms, such as deubiquitination and phosphorylation (25, 26). First, E7 binds to several ubiquitin-specific proteases (USPs) that remove ubiquitin from ubiquitinated proteins (47). Previous studies have shown that USP7 and USP11 deubiquitinate and stabilize E7 proteins (25, 48). In addition, dual-specificity tyrosine phosphorylation-regulated kinase 1A (DYRK1A) phosphorylates HPV16 E7 and prolongs the E7 protein half-life by hindering ubiquitinated E7 binding to proteasomes (26). E7 proteins are also stabilized by an E6^E7 fusion protein from an E6^E7 splice isoform that encodes 41 N-terminal amino acids of E6 and 38 C-terminal amino acids of E7 (49). The E6^E7 fusion protein binds to and stabilizes both E6 and E7 proteins in the presence of GRP78 and HSP90 (49). A recent study has also shown that E6AP overexpression stabilizes E7 protein in a proteasome-dependent manner (50). While these previous reports suggest that E7 protein can be stabilized through various mechanisms, it is still unknown how E7 protein overcomes its degradation mediated by the SCF ubiquitin ligase complex containing CUL1 and UBE2L3. Our findings reveal a novel mechanism of E7 protein stabilization by degrading CUL1 and UBE2L3, the important components of the SCF ubiquitin ligase complex.

The SCF ubiquitin ligase complex is characterized by a substrate recognition factor for connecting the substrate to the E3 ubiquitin ligase complex. The adaptor is a limiting factor for the targeted process and substrates. The SCF ubiquitin ligase complex has a central role in cell-cycle regulation. Deregulated cell-cycle control is a hallmark of cancer. The F box protein adaptor targets negative cell cycle regulators, such as cyclins (A, D, and E) (51), p21 (18), p27 (19), and p57 (52), for ubiquitination and degradation (53). Hence, it promotes cell cycle progression during S and G2 phases (53). Several studies have shown that CUL1 upregulation and its association with poor patient prognosis in various cancers, including gastric, breast, renal cell carcinoma, colorectal, and hepatocellular carcinoma (54–58). Additionally, CUL1 augments cancer cell proliferation, adhesion, migration, and metastasis (54–59). On the other hand, the SCF ubiquitin ligase complex binds to RNA-binding proteins of the fragile X protein family FMRP, FXR1, and FXR2 in HPV-negative HNC cells (60). The binding of the SCF ubiquitin ligase complex to these RNA-binding proteins led to their degradation and suppressed tumorigenesis (60). Additionally, our results show that CUL1 overexpression in HPV+ HNC cells hinders cell proliferation and delays tumor growth in vivo.

Ubiquitination and degradation of p53 by E6 and E6AP are considered hallmarks of HPV-associated cancer progression. UBE2L3, an E2 conjugating enzyme, is essential in the E6AP E3 ubiquitin ligase complex for ubiquitinating p53 (61). However, UBE2L3 upregulation by AhR activation in HeLa cells inhibits cell proliferation and enhances apoptosis (62, 63). This result may be caused by E7 degradation as UBE2L3 and CUL1 ubiquitinate and degrade E7 proteins in cervical cancer cells (17). In contrast, UBE2L3, associated with the SCF ubiquitin ligase complex, promotes non-small-cell lung cancer by ubiquitinating and degrading p27kip1 and GSK-3β (64, 65). Thus, CUL1 and UBE2L3 might have double-edged functions for tumor promotion in lung cancer while suppressing HPV+ cancers through E7 degradation.

We report here that MARCHF8 plays an important role in E7 protein stabilization in HPV+ HNC cells by degrading CUL1 and UBE2L3 (**Fig. 8**). As E7 is required for viral replication, this finding suggests that MARCHF8 contributes to HPV infection and persistence. Likewise, recent studies have shown that MARCHF8 supports infection of several human viruses, including HCV, dengue, and Zika (66). Particularly, MARCHF8 induces the polyubiquitination of the HCV nonstructural2 (NS2) protein that mediates binding to ESCRT0 and assembling viral envelope (67, 68). Additionally, MARCHF8 facilitates HIV infection by ubiquitinating and degrading Tetherin independent of Vpu (69). In contrast, MARCHF8 is also known as an antiviral restriction factor (66). MARCHF8 targets several viral envelope glycoproteins to inhibit infections of HIV, vesicular stomatitis virus, rabies virus, lymphocytic choriomeningitis virus, SARS-CoV, Chikungunya virus, and Ross River virus (70–72). Moreover, MARCHF8 also restricts Influenza A virus infectivity by inhibiting the incorporation of the virus glycoproteins into the progeny virions (73) and targeting the M2 protein for ubiquitin-dependent degradation (74). These findings suggest that MARCHF8 plays a dual role in virus infection as a proviral and antiviral factor.

**Fig. 8.**
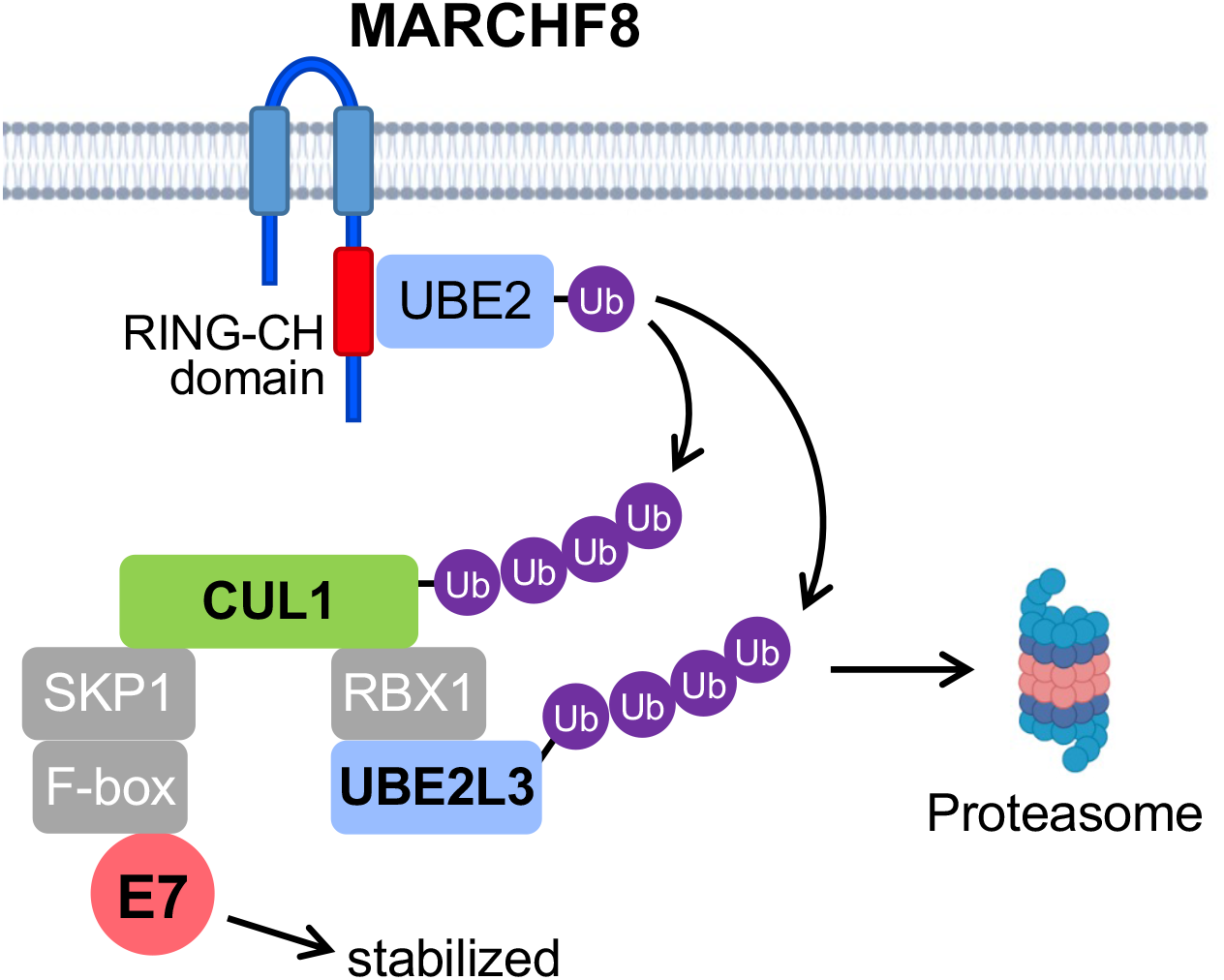
The schematic models of MARCHF8-mediated E7 stabilization by degrading CUL1 and UBE2L3. The HPV oncoprotein E6 activates the *MARCHF8* promoter activity through the MYC/MAX transcription factor complex and upregulates MARCHF8 in the HPV+ HNC cells (31). MARCHF8 protein ubiquitinates and degrades CUL1 and UBE2L3 proteins, leading to the prevention of E7 protein degradation.

As E7 is required for HPV replication and carcinogenesis, destabilizing E7 proteins by targeting MARCHF8 may effectively prevent and treat HPV-associated cancers. Additionally, we have recently shown that MARCHF8 ubiquitinates and degrades death receptors to impede apoptosis of HPV+ HNC cells and that *MARCHF8* knockout significantly suppresses tumor growth in vivo (31). Taken together, our findings suggest MARCHF8 as a potential therapeutic target for HPV+ HNC.

## Materials and Methods

**Cell lines:** 293FT cells were obtained from Thermo Fisher (Waltham, MA). HPV+ HNC (SCC2, SCC90, and SCC152) and HPV-HNC (SCC1, SCC9, and SCC19) cells were purchased from the American Type Culture Collection (Manassas, VA). These cells were maintained in Dulbecco’s modified Eagle’s medium (DMEM) containing 10% fetal bovine serum (FBS) and penicillin/streptomycin (Thermo Fisher) as described (12, 75–77). The N/Tert-1 cell (78) expressing HPV16 E6 (N/Tert-1-E6), E7 (N/Tert-1-E7), and E6 and E7 (N/Tert-1-E6E7) were previously established (79, 80) and cultured in keratinocyte serum-free medium containing epidermal growth factor (EGF), bovine pituitary extract, and penicillin/streptomycin (Thermo Fisher). The mouse oropharyngeal epithelial (MOE) cell lines, normal immortalized mouse oral epithelial cell (NiMOE), mEERL (HPV+), and MOE/shPtpn13 (HPV-), were obtained from John Lee (34) and maintained in E-medium (DMEM and F12 media containing 0.005% hydrocortisone, 0.05% transferrin, 0.05% insulin, 0.0014% triiodothyronine, 0.005% EGF, and 2% FBS) as previously described (81).

### Lentivirus constructs and production

The human *CUL1* and *UBE2L3* coding sequences were amplified from pcDNA-HA-UBE2L3 (Addgene, #27561) and pcDNA-myc3-CUL1 (Addgene, #19896), respectively and cloned into the pLenti6/V5-D-TOPO backbone (Addgene, #22945). The mouse *Cul1* and *Ube2l3* coding sequences were amplified from cDNA prepared from NiMOE cells and cloned into the pLenti6/V5-D-TOPO backbone (Addgene, #22945). The primer sequences used in amplifying human and mouse *CUL1* and *UBE2L3* coding sequences are listed in **Table 1**. The shRNAs targeting human *MARCHF8* were ordered from Sigma-Aldrich (St. Louis, MO). The sgRNAs targeting *Marchf8* were designed using the web-based software ChopChop (<u>chopchop.cbu.uib.no</u>) (82) and cloned into the lentiCRISPR v2-blast plasmid (Addgene, #83480) using ligating duplex oligonucleotides containing BsmBI restriction sites purchased from Integrated DNA Technologies (IDT, Coralville, IA). The shRNAs and sgRNAs used are listed in **Tables 2 and 3**. Lentiviruses containing human and mouse *CUL1*, *UBE2L3*, and shRNA or sgRNA against MARCHF8 were produced using 293FT cells with packaging constructs, pCMV-VSV-G (Addgene, #8454) and pCMV-Delta R8.2 (Addgene, #12263). The lentiviruses were harvested 48 hrs post-transfection and concentrated by ultracentrifugation at 25,000 rpm for 2 hrs. In addition, cells were incubated with lentiviruses for 48 hrs in the presence of polybrene (8 μg/ml) and selected with blasticidin (8 μg/ml).

**Table 1.**
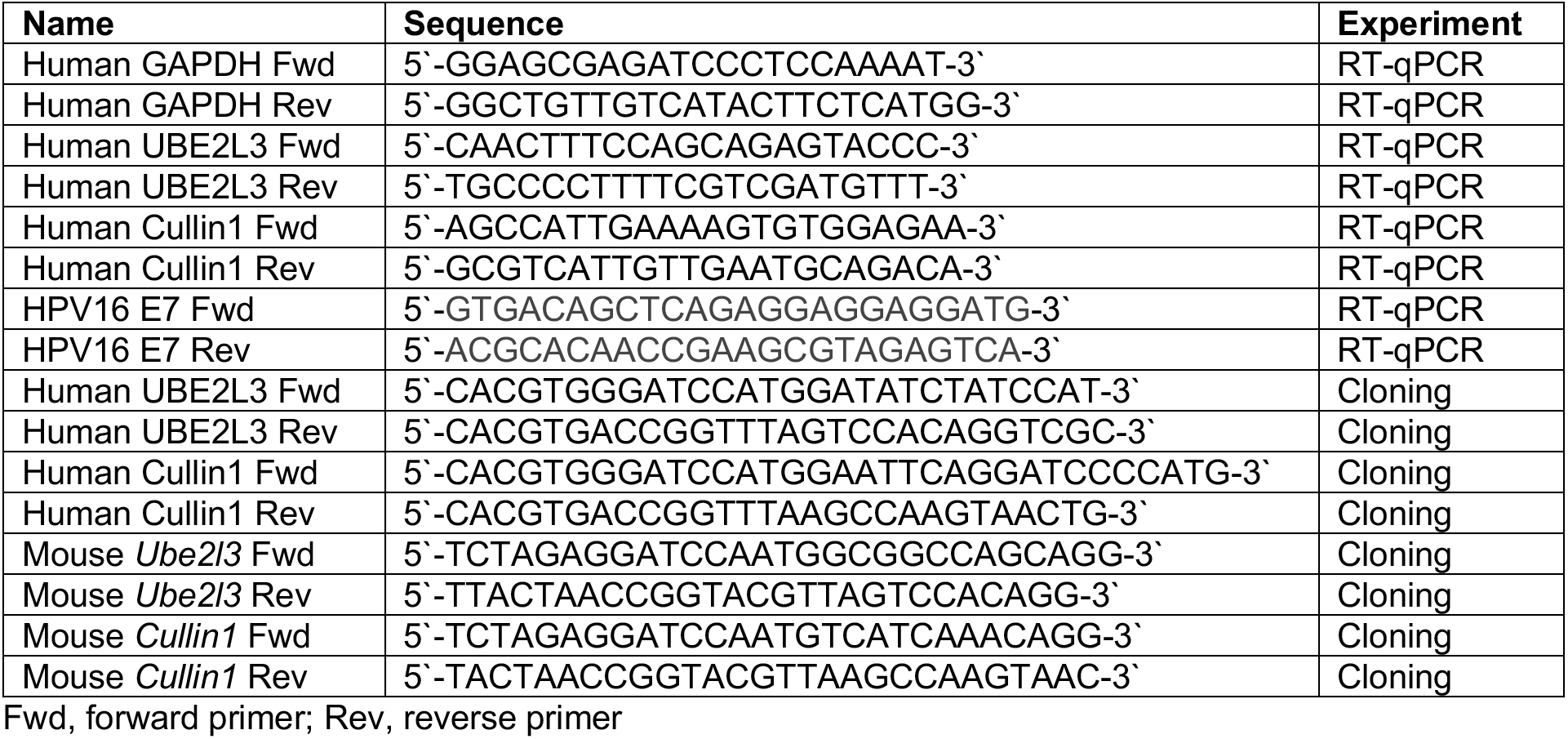
List of the oligonucleotides.

**Table 2.**
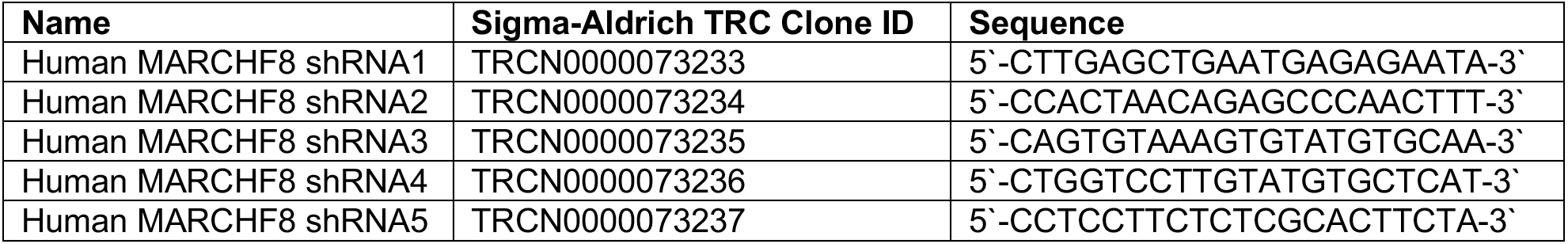
List of the shRNAs.

**Table 3.**
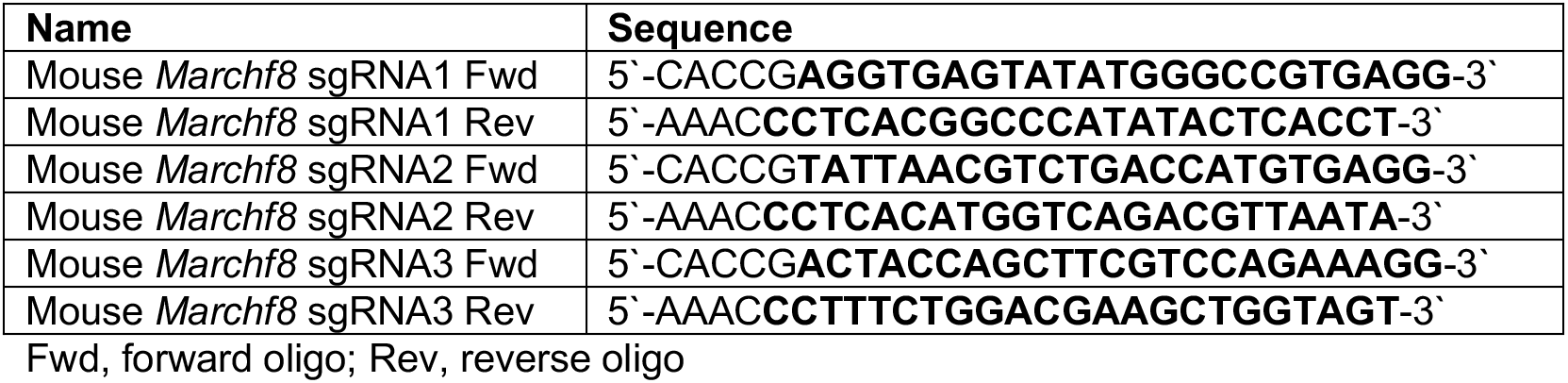
List of the sgRNAs.

### Bromodeoxyuridine (BrdU) staining

1 × 10^6^ SCC152 and SCC2 MARCHF8 knockdown and scrambled cells were seeded per 10 cm^2^ petri dish in DMEM supplemented with 10% FBS and penicillin/streptomycin. Cells were incubated with a medium containing 10 mM BrdU for 16 hours. Cells were trypsinized and stained using a BrdU staining kit (Thermo Fisher) according to the manufacturer’s instructions. Cell viability was determined by staining cells using the Zombie NIR kit (BioLegened). Stained cells were analyzed using an LSRII flow cytometer.

### Immunoprecipitation (IP) and western blotting

Whole cell lysates were prepared in 1X radioimmunoprecipitation assay buffer (RIPA) buffer (Abcam, Waltham, MA) containing protease inhibitor cocktail (Roche, Mannheim, Germany) according to the manufacturer’s instructions. The protein concentration was measured by the Pierce BCA Protein Assay Kit (Thermo Fisher). IP was performed using the Pierce Classic Magnetic IP/Co-IP Kit (Thermo Fisher). Briefly, 25 μl of protein A/G magnetic beads were incubated with 5 μg of specific antibodies (**Table 4**) for two hours. 1-2 mg of the whole cell lysates were incubated with antibody-coupled beads overnight at 4°C. In addition, 10-20 μg of protein was used to perform western blotting with antibodies listed in **Table 4** as previously described (83). NIH ImageJ software was used to determine band intensities and normalized to the β-actin band intensity.

**Table 4.**
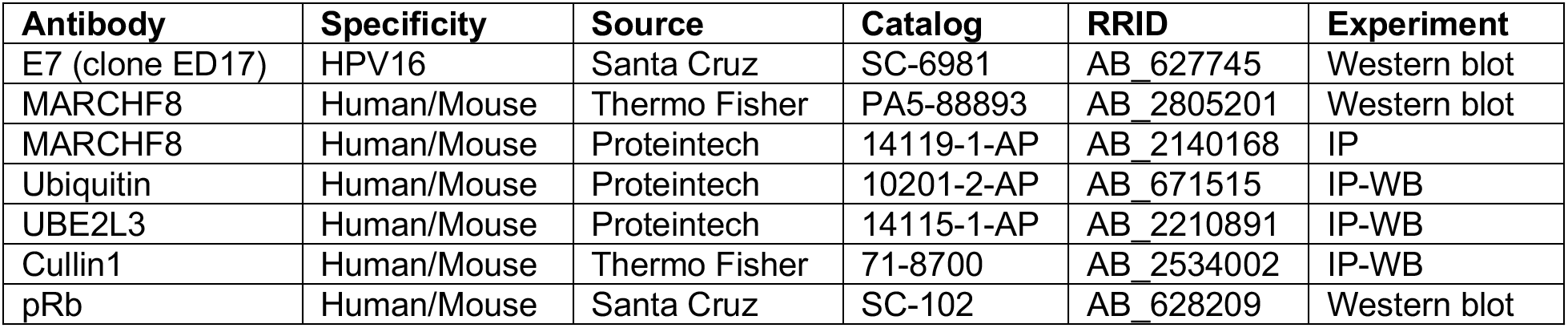
List of the antibodies.

| Antibody | Specificity | Source | Catalog | RRID | Experiment |
| --- | --- | --- | --- | --- | --- |
| E7 (clone ED17) | HPV16 | Santa Cruz | SC-6981 | AB_627745 | Western blot |
| MARCHF8 | Human/Mouse | Thermo Fisher | PA5-88893 | AB_2805201 | Western blot |
| MARCHF8 | Human/Mouse | Proteintech | 14119-1-AP | AB_2140168 | IP |
| Ubiquitin | Human/Mouse | Proteintech | 10201-2-AP | AB_671515 | IP-WB |
| UBE2L3 | Human/Mouse | Proteintech | 14115-1-AP | AB_2210891 | IP-WB |
| Cullin1 | Human/Mouse | Thermo Fisher | 71-8700 | AB_2534002 | IP-WB |
| pRb | Human/Mouse | Santa Cruz | SC-102 | AB_628209 | Western blot |

### Reverse transcription-quantitative PCR (RT-qPCR)

RNeasy Plus Mini Kit (Qiagen, Germantown, MD) was used to extract total RNA. First-strand cDNA was synthesized from 2 μg of total RNA using Superscript II reverse transcriptase (Roche, Mannheim, Germany). qPCR was performed in a 20 μl reaction mixture containing 10 μl of SYBR Green Master Mix (Applied Biosystems), 5 μl of 1 mM primers, and 100 ng of cDNA template using a Bio-Rad CFT Connect thermocycler. Data were normalized to glyceraldehyde 3-phosphate dehydrogenase (GAPDH). IDT synthesized primers used in qPCR can be found in Supplementary **Table 1**.

### Mice and tumor growth

C57BL/6J mice were purchased from Jackson Laboratory (Bar Harbor, ME) and maintained by the USDA guidelines. 5 × 10^5^ mEERL cells were subcutaneously injected into the rear right flank of 6 to 8-week-old mice (*n* = 10 per group). Tumor volume was measured twice weekly and calculated using the equation: volume = (width^2^ X length)/2. Mice were euthanized when tumor volume reached 2000 mm^3^, as previously described (84). Animals were determined as tumor-free when no measurable tumor was observed 12 weeks post-injection. Survival curves were generated by the Kaplan-Meier method with a tumor volume of 2000 mm^3^ as an endpoint. The Michigan State University Institutional Animal Care and Use Committee (IACUC) approved experiments by National Institutes of Health guidelines for using live animals.

### Statistical analysis

Data were analyzed using GraphPad Prism (San Diego, CA) and were presented as mean ± standard deviation. Statistical significance was determined using an unpaired Student’s *t-test*. *P* values <0.05 are considered statistically significant. Distributions of time-to-event outcomes (e.g., survival time) were summarized with Kaplan–Meier curves and compared across groups using the log-rank test with α = 0.01.

## ACKNOWLEDGMENT

We thank members of the Pyeon laboratory for their valuable comments and suggestions. This work was supported by NIH R01 DE026125 and R01 DE029524 (DP) and the Michigan State University Global Impact Initiative (DP).

